# Priors, population sizes, and power in genome-wide hypothesis tests

**DOI:** 10.1101/737676

**Authors:** Jitong Cai, Jianan Zhan, Dan E. Arking, Joel S. Bader

**Author notes:** 23andMe, Inc (Jianan Zhan).

## Abstract

Genome-wide tests, including genome-wide association studies (GWAS) of germ-line genetic variants, driver tests of cancer somatic mutations, and transcriptome-wide association tests of RNA-Seq data, carry a high multiple testing burden. This burden can be overcome by enrolling larger cohorts or alleviated by using prior biological knowledge to favor some hypotheses over others. Here we compare these two methods in terms of their abilities to boost the power of hypothesis testing. We provide a quantitative estimate for progress in cohort sizes, and present a theoretical analysis of the power of oracular hard priors: priors that select a subset of hypotheses for testing, with an oracular guarantee that all true positives are within the tested subset. This theory demonstrates that for GWAS, strong priors that limit testing to 100–1000 genes provide less power than typical annual 20–40% increases in cohort sizes. These theoretical results explain the continued dominance of simple, unbiased univariate hypothesis tests for RNA-Seq studies and GWAS: if a statistical question can be answered by larger cohort sizes, it should be answered by larger cohort sizes rather than by more complicated biased methods involving priors. We suggest that priors are better suited for non-statistical aspects of biology, such as pathway structure and causality, that are not yet easily captured by standard hypothesis tests.

**Author summary:** Biological experiments often test thousands to millions of hypotheses. Gene-based tests for human RNA-Seq data, for example, involve approximately 20,000 tests; genome-wide association studies (GWAS) involve about 1 million effective tests. A robust approach is to perform individual tests and then apply a Bonferroni correction to account for multiple testing. This approach implies a single-test p-value of 2.5 × 10^−6^ for RNA-Seq experiments, and a p-value of 5 × 10^−8^ for GWAS, to control the false-positive rate at a conventional value of 0.05. Many methods have been proposed to alleviate the multiple-testing burden by incorporating a prior probability that boosts the significance for a subset of candidate genes or variants. At the extreme limit, only hypotheses within a candidate set are tested, corresponding to a decreased multiple testing burden. Despite decades of methods development, prior-based tests have not been generally used. Here we compare the power increase possible with a prior with the power increase from a much simpler strategy of increasing a study size. We show that increasing the population size is exponentially more valuable than increasing the strength of prior, even when the true prior is known exactly. Furthermore, even modest yearly increases in actual GWAS cohorts can yield power gains beyond the reach of any reasonable prior. These results provide a rigorous explanation for the continued use of simple, robust methods rather than more sophisticated approaches. They suggest that the value of priors is not in multiple hypothesis testing but rather in non-statistical aspects of interpretation including pathway structure and causality.

## Introduction

Genomics experiments involve testing thousands to millions of hypotheses. In functional genomics and proteomics, each gene or protein usually corresponds to a single test, with 20,000 or more tests required for an RNA-Seq or proteomics experiment. In human genetics, the number of independent tests accounting for linkage disequilibrium in a single ethnicity is usually assumed to be about 1 million for all but the rarest variants. To maintain a family-wise error rate (FWER) controlled at 0.05, a long-standing approach has been to apply a Bonferroni correction, requiring a single-test p-value of 0.05 divided by the number of hypotheses tested. This multiple-testing correction from this stringent approach is a burden for identifying genome-wide significant findings.

A robust solution to this problem has been to gather large cohorts of unrelated individuals, particularly for GWAS [1]. While the biological effect of a genetic variant is constant, its corresponding test statistic should be improved with cohort size, yielding greater power to detect. Cohort sizes are limited by the efficiencies of data generation, for example the number of samples that can be genotyped for a typical research budget. Increased DNA sequencing efficiencies permit larger cohorts for RNA-Seq, and increased DNA synthesis efficiencies reduce the cost of genotyping arrays and permit larger cohorts. Progress in exponentially improving fields is often characterized by the doubling time, popularly known as Moore’s law for 1.5–2 year doubling time for the number of transistors on a semiconductor computer chip [2]. Moore’s law analysis applied to the number of DNA bases that can be sequenced or synthesized per dollar has shown a doubling time of approximately 2 years [3]. Of course, sequencing or genotyping costs are only one aspect of a study, and actual cohort sizes may grow at different rates.

Rather than increasing cohort sizes, an alternative approach is to incorporate prior knowledge about functional effects of genes or SNPs. In GWAS, this may increase the power to detect SNPs with true associations or to identify which SNP in a linkage disequilibrium (LD) region is most likely to be the causal variant [4–7]. Other methods incorporate priors based on patterns learned from the data, for example priors for gene-based patterns [8,9] or phenotype-based patterns [10]. While these methods have value in providing a clearer view of genetic architecture than available through univariate tests, the number of new significant findings has been small [7,11].

A representative approach incorporated 450 different annotations into GWAS analysis of 18 human traits; the number of loci with high-confidence associations was increased by around 5% [12]. Despite the intuitive value of incorporating pre-existing biological knowledge, it remains unclear whether this roughly 5% increase in genome-wide significant findings is the best that could be obtained, and additionally whether the increase comes at the cost of false negatives for true positives that lack similar annotations. It is also unclear how this 5% increase compares with the anticipated increase from cohort size alone: given that this more sophisticated analysis itself required 1-2 years of effort, would it have been just as effective to wait a year and then apply simpler methods to a larger cohort?

In this paper, we use theoretical models and derivations to investigate the dependency of power on population size and incorporating priors. We consider an oracular hard prior, which tests a subset of the hypothesis that is guaranteed to include all the true positives. We show analytically that in the limit of small effect sizes and most relevant to genomics studies, population sizes are exponentially more important than priors in determining the power. We then show that given historical trends in cohort sizes, it is nearly impossible for new analytical methods to improve power faster than larger studies that use conventional methods.

## Methods

### Empirical data

Data sets were collected from the GWAS Catalog (https://www.ebi.ac.uk/gwas/, accessed on June 13, 2020) [13]. An effective sample size was calculated for each study. For studies investigating quantitative traits, the effective sample size was estimated as the number of individuals in the largest cohort described. For case-control studies, the effective sample size was calculated as twice the harmonic mean of the case and control population, estimated to have equal power as follows.

Consider a study with *N*_1_ cases and *N*_2_ controls. In the context of GWAS, tests for each allele are typically based on *δp*, the difference in allele frequency between cases and controls. Denoting these allele frequencies as *p*_1_ and *p*_2_,

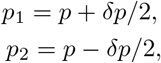

The test statistic is *Q*^2^,

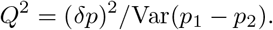

Under the null hypothesis, *δp* = 0 and *Q*^2^ follows a *χ*^2^ distribution for one degree of freedom.

The estimated allele frequency 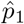 is the observed allele count, 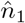, divided by the number of chromosomes, 2*N*_1_. The allele count itself is a binomial random variable with expectation 2*N*_1_*p*_1_ and variance 2*N*_1_*p*_1_(1 – *p*_1_). The variance of *p*_1_ is therefore

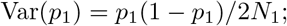

similarly,

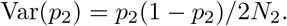

Under the null hypothesis, and for small effect sizes that are typical in GWAS, *δp* is small. Neglecting terms of order *δp*,

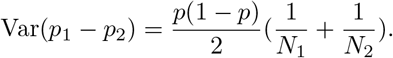

Now suppose that a study involves a population with total size *N*. The variance is minimized with *N*_1_ = *N*_2_ = *N*/2, suggesting our definition of effective population size *N* according to

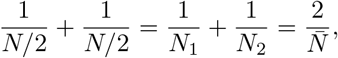

where 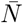 is the harmonic mean 2*N*_1_*N*_2_/(*N*_1_ + *N*_2_). For the effective population size,

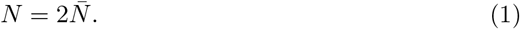

Each GWAS study may be related to multiple records, each of which demonstrates the significant association between the trait investigated by the study and a SNP. These records are generally account for linkage disequilibrium by reporting the most significant SNP in a linkage region. The number of associations for each study was counted as the number of associations passing the genome-wide significance threshold of 5 × 10^−8^.

Studies were grouped into phenotypes based on the trait vocabulary. For every phenotype, we then arranged the studies chronologically by publication date and retained studies reporting more findings than all previous studies. Only phenotypes with at least 3 effective studies were kept for further analysis.

Doubling times for cohort sizes and number of associations for each phenotype were estimated as a linear model, log_2_ *ŷ* = *β*_0_ + *β*_1_*t*, with *y* representing either the cohort size or the number of associations, *t* the publication date with months and days converted to fractional years, and regression coefficients *β*_0_ and *β*_1_, corresponding to exponential growth, *ŷ* = 2^*β*_0_2^ 2^*β*_1_*t*^. The doubling time is *τ* ≡ 1/*β*_1_, and its error is 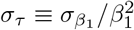.

Significance of the exponential fit was assessed by three nested regression models denoted *R*_0_, *R*_1_, and *R*_2_: the null model of no progress, *R*_0_: log_2_ *ŷ* = *β*_0_; the log-linear model of exponential growth, *R*_1_: log_2_ *y* = *β*_0_ + *β*_1_*t*; and a model with more complicated quadratic time dependence, *R*_2_: log_2_ *y* = *β*_0_ + *β*_1_*t* + *β*_2_*t*^2^. Significance of the exponential growth model relative to a null hypothesis of no progress was estimated as the ANOVA p-value of *R*_1_ versus *R*_0_. For traits with significant growth, we then assessed the evidence for more complicated time dependence as the ANOVA p-value of *R*_2_ versus *R*_1_. Note that for a specific trait, large cohort studies could in principle identify all loci for that trait, yielding a significant model for cohort size growth but no growth in the number of significant loci. The F-statistic of ANOVA test for model sufficiency was calculated as the proportion of extra variation explained by the full model compared to the reduced model. Specifically, to test the time dependence of growth, F-statistic was calculated as the ratio of extra variation explained by *R*_1_ compared to *R*_0_ and the variation explained by *R*_1_ alone; to test whether growth had a more complicated quadratic time dependence, F-statistic was calculated as the ratio of extra variation explained by *R*_2_ compared to *R*_1_ and the variation explained merely by *R*_2_. The corresponding p-value was then calculated by the F-statistic.

Doubling times for cohort size (*τ_N_* and error *σ_N_* calculated as described above) and number of significant loci (*τ_L_* and *σ_L_* calculated as described above) were compared for a test of the null hypothesis that *τ_N_* = *τ_L_*. The test statistic *z_τ_* was defined as 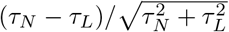, and p-values were calculated for a two-sided test of the null hypothesis *z_τ_* = 0. As this analysis was exploratory, we did not correct this test for the number of traits analyzed.

### Relating the significance threshold, power, cohort size, and variance explained for genome-wide tests

We consider tests of association between a feature of the data, *x*, and an observed phenotype or response variable, *y*, assumed to be scalars for simplicity. For a population of size *N*, these are aggregated into vectors **x** and **y**. An association test compares a null model *M*_0_, to an alternative, *M*_1_, which for a linear model takes the form

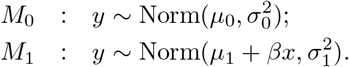

One such *M*_1_ exists for each possible feature to be tested. With *A* total possible alternatives to be tested, these could be denoted {*M_a_*}, *a* ∈ {1, 2,…, *A*}. We consider one such alternative at a time and for simplicity denote it *M*_1_. Model parameters are 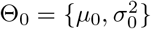 for the null model and 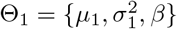 for the alternative model. These models correspond to a null hypothesis *H*_0_ and alternative hypothesis *H*_1_,

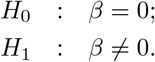

For nested models, the hypothesis test is usually performed by a likelihood ratio test or its equivalent. Denote the maximum likelihood parameters as 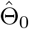 and 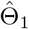, and assume independence of the model and data. A test statistic *τ* is defined as

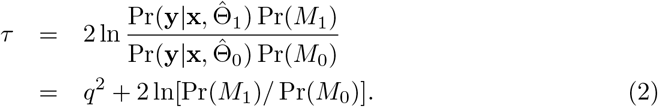

According to Wilks’ Theorem, under the null hypothesis, *q*^2^ is a random variable distributed as 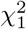, or more generally as a 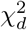, random variable where the null model is nested inside an alternative model with *d* additional parameters [14]. Under the alternative hypothesis, *q*^2^ is distributed as a non-central *χ*^2^ with non-centrality parameter 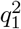,

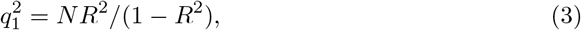

where *R*^2^ is the fraction of variance explained by the alternative hypothesis, and 1 – *R*^2^ is the residual fraction of variance.

For a conventional test, the prior Pr(*M*) is identical for the null and each alternative; it does not contribute to the test statistic. To control the type I error (false-positive rate) at family-wise error rate FWER *α*, the Bonferroni method requires a single-test p-value of *α/A* for *A* total tests. Let the quantile of the uniform normal distribution corresponding to a two-tailed test at this stringency be *z_I_*. More formally, if Φ(*z*) is the cumulative lower tail probability distribution for standard normal random variable *z*, then Φ(−*z_I_*) = *α*/2*A*. For true effect *q*_1_, the power is Φ(|*q*_1_| – *z_I_*), or equivalently

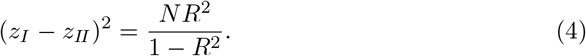

This key expression relates the type I error (false-positive rate), the type II error (false-negative rate or complement of power), the population size *N*, and the effect size *R*^2^.

### Effect size distribution

In the limit of small effect size, *R*^2^ ≪ 1, and fixed type I and type II error, the effect size and population size are inversely related,

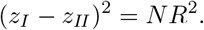

This relationship, together with doubling times, implies a functional form for the number of loci with effect size *R*^2^ or larger, defined as *L*(*R*^2^). As before, define *τ_N_* as the doubling time for cohorts and *τ_L_* as the doubling time for loci. The effect size that can be discovered at a specified type I error is approximately equal to the effect size at 50% power,

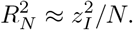

After *t* years, cohort size increases from an initial value *N*_0_ to a final value *N_t_* = *N*_0_2^*t*/*τ_N_*^. Similarly, the number of loci discovered increases from *L*_0_ to *L*_0_2^*t/τ_L_*^.

The number of loci at the end is also equal to the the number of loci with effect size greater than or equal to 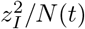,

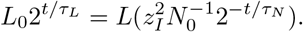

This relationship is satisfied in turn by a power-law dependence of *L*(*R*^2^) on its argument,

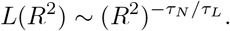

The probability density has the form of the derivative of the cumulative probability, and thus well-defined doubling times imply an effect-size probability distribution *ρ*(*R*^2^) with functional form

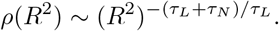

### Oracular hard priors

We consider an idealized prior in which only hypotheses corresponding to a faction 1/*S* of the total are tested, with an oracular property that all known positives lie within the selected subset. Larger *S* corresponds to a stronger prior. For 20,000 gene-based tests, testing 10% of the total corresponds to *S* = 10, and testing 20 genes corresponds to *S* = 1000. Realistically, priors stronger than *S* = 100, corresponding to 200 genes tested, are unlikely.

The effect of a hard prior is to reduce the multiple-testing burden. To maintain FWER *α*, each two-tailed test is performed at stringency *Sα*/2*A* rather than *α*/2*A*. This reduces the quantile *z_I_* required for significance and increases the power to detect an association with a smaller effect *R*^2^. Equivalently, Eq. 4 can be solved for *R*^2^ to calculate the critical effect size to achieve desired power at stated type I error,

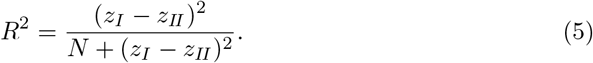

The effect of a hard prior on *z_I_* may also be estimated analytically. A steepest descents approximation relates the quantile *z* > 0 to its upper-tail area *ϵ*,

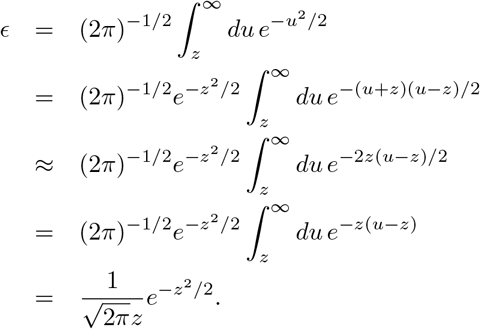

Equivalently,

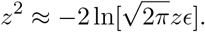

In terms of the quantile *z_I_* for prior strength *S* and a two-tailed test, we have approximately

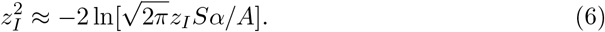

Define *ζ* as the value of *z_I_* for no prior, *S* =1, with Φ(−*ζ*) = *α*/2*A* and

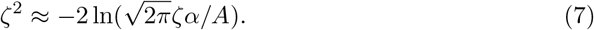

For GWAS with a p-value threshold of 5 × 10^−8^, *ζ* = 5.45 and *ζ*^2^ = 29.7. Because the dependence of Eq. 6 on ln *z* is weak, we replace ln *z* with ln *ζ*,

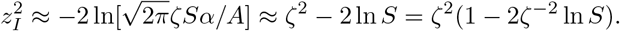

Keeping terms of order 1/*ζ*,

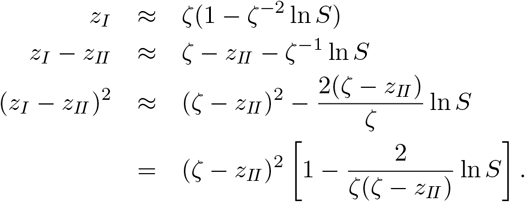

According to Eq. 4, the critical effect size depends only on the ratio (*z_I_ – z_II_*)^2^/*N*. Consider two scenarios with equal critical effect size, one with population size *N*_1_ and prior strength *S*_1_, and the second with population size *N*_2_ and prior strength *S*_2_. For these to have equal critical effect size,

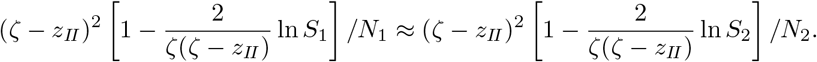

Cancelling constant terms *ζ* – *z_II_* and noting that 2*ζ*^−1^(*ζ* – *z_II_*) ln *S* is small,

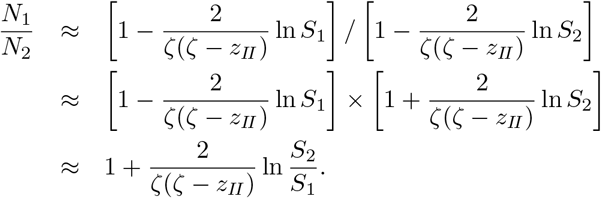

The dependence on population size is linear, whereas the dependence on prior strength is logarithmic. Equivalently, population size is exponentially more important that prior strength. Again for GWAS with *z_II_* selected for 80% power, *ζ*(*ζ* – *z_II_*)/2 = 17.15, and only a small fractional population increase is required to obtain the equivalent power increase for a strong prior. In the equation above, the logarithmic term divided by *ζ*(*ζ* – *z_II_*)/2 is small, permitting the approximation 1 + *ϵ* ≈ *e^ϵ^* with error *ϵ*^2^. Using this approximation,

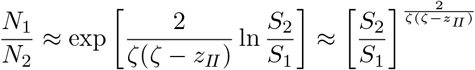

An extremely strong prior with *S*_2_ = 1000, with effectively only 20 genes selected for testing, can be matched by a population increase of about 40%.

Contours of *N* and *S* with equal critical effect size can be estimated by returning to the approximate result

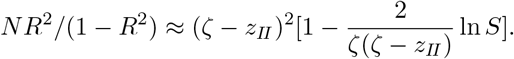

Noting that for small *ϵ*, 1 + *ϵ* ln *S* ≈ *S^ϵ^*, contours are given by

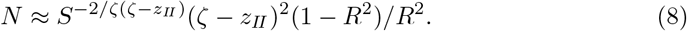

On a log-log plot of log *S* versus log *N*, these contours would have steep negative slope equal to −*ζ*(*ζ* – *z_II_*)/2.

## Results

### Doubling times for GWAS cohorts and significant loci

Exponential increases in genotyping and sequencing efficiency have enabled similar increases in GWAS cohort sizes. Larger sample sizes in turn have greater power to detect SNPs with smaller and smaller effects. We quantified this relationship through a systematic analysis of GWAS cohorts, traits, and loci as compiled by the GWAS catalog [13].

GWAS catalog contains results for 5,123 total studies describing 3,034 traits and 126,788 associations that are genome-wide significant with p-value 5 × 10^−8^ or below. Studies were grouped according to the catalog-assigned disease trait and arranged in chronological order from the oldest to the most recent. Cohort sizes were based on the population reported by the study. Case-control population sizes were estimated as twice the harmonic mean of the number of cases and controls, a balanced design that should have similar power (see Methods). To avoid analysis of smaller replication studies, studies for a trait were only analyzed if the study identified the largest number of significant loci for that trait as of its publication date. Traits with at least three increasing number of findings were then fit to a regression model to estimate the doubling time for cohort size and for number of significant loci.

Results for a well-studied case-control disease trait, breast cancer susceptibility, demonstrate the progress in identifying genome-significant loci (Fig. 1). As of 2010, the largest study had 3,659 cases and 4,897 controls, an effective cohort size of 8,377. Those studies had revealed 19 genome-wide significant loci. As of 2015, the largest effective cohort size was 33,671, and 106 loci had been identified. Studies through 2020 have had an effective cohort size of 138,040 and 411 loci have been identified.

**Fig 1.**
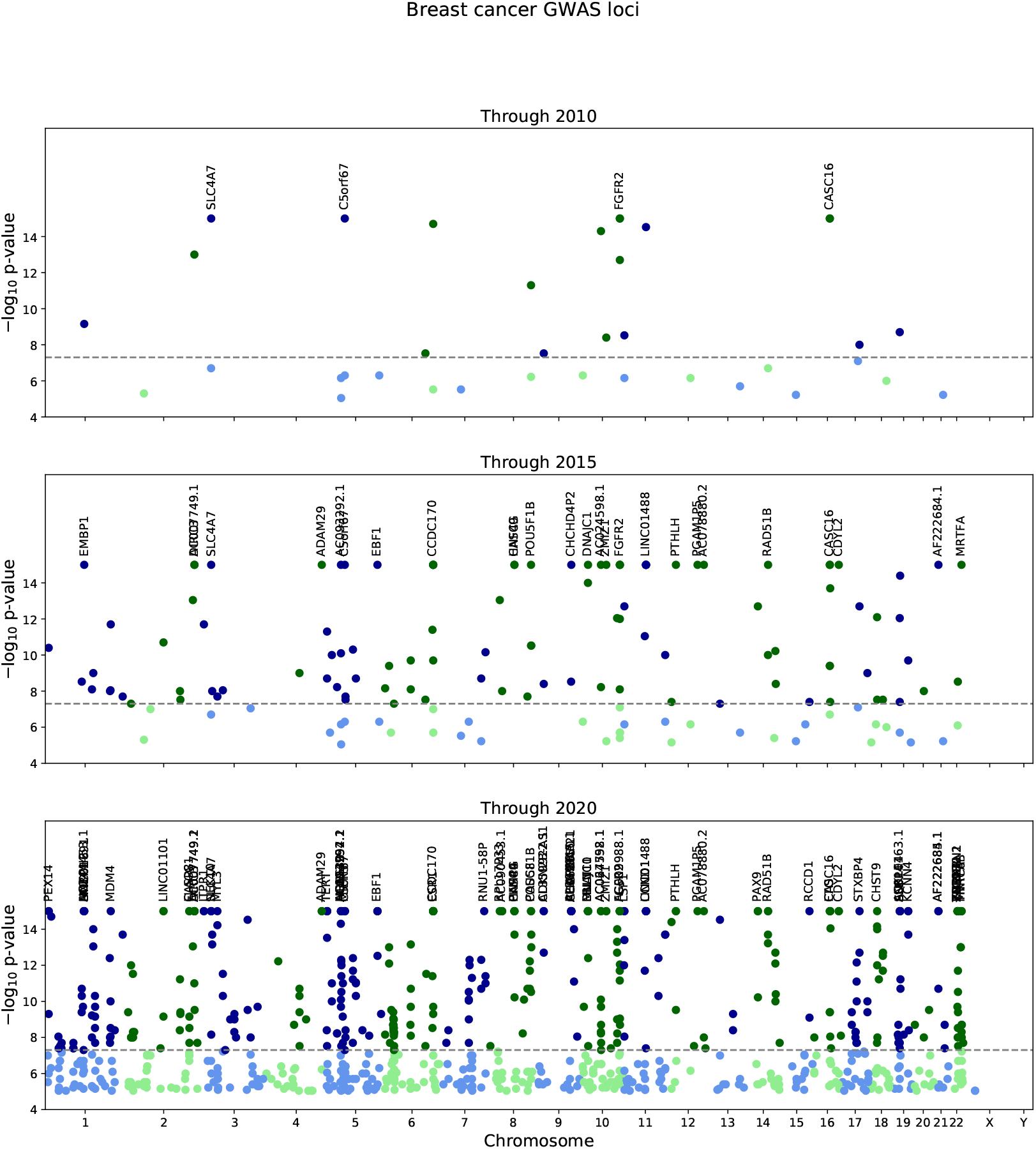
GWAS progress for breast cancer susceptibility cohorts and loci. Manhattan plots depict GWAS findings for breast cancer as of 2010 (top panel), 2015 (middle panel), and 2020 (bottom panel). In each panel, the x-axis represents genomic coordinates to scale, and the y-axis is the −log_10_ p-value for a GWAS association with a SNP; a dashed line indicates the genome-wide significance threshold, *p* = 5 × 10^−8^. The SNP color alternates blue/green by chromosome, with lighter colors for findings below threshold and saturated colors above threshold.

Regression fits for the five breast cancer susceptibility studies estimate the cohort size doubling time to be *τ_N_* = 1.5 ± 0.2 years, and the genome-wide significant loci doubling time to be *τ_L_* = 1.6 ± 0.2 years (Fig. 2). This fit, highly significant versus a null hypothesis of no progress, used the log-linear model log_2_ *y* = *β*_0_ + *β*_1_*t*, where *y* is the cohort size or the number of genome-wide significant loci, *t* is the publication date in fractional years, and 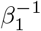 is the doubling time. A fit including a quadratic term, log_2_ *y* ~ *β*_0_ + *β*_1_*t* + *β*_2_*t*^2^, did not improve the model for cohort doubling time (p = 0.384) or for loci doubling time (p = 0.652), justifying the use of an exponential growth model.

**Fig 2.**
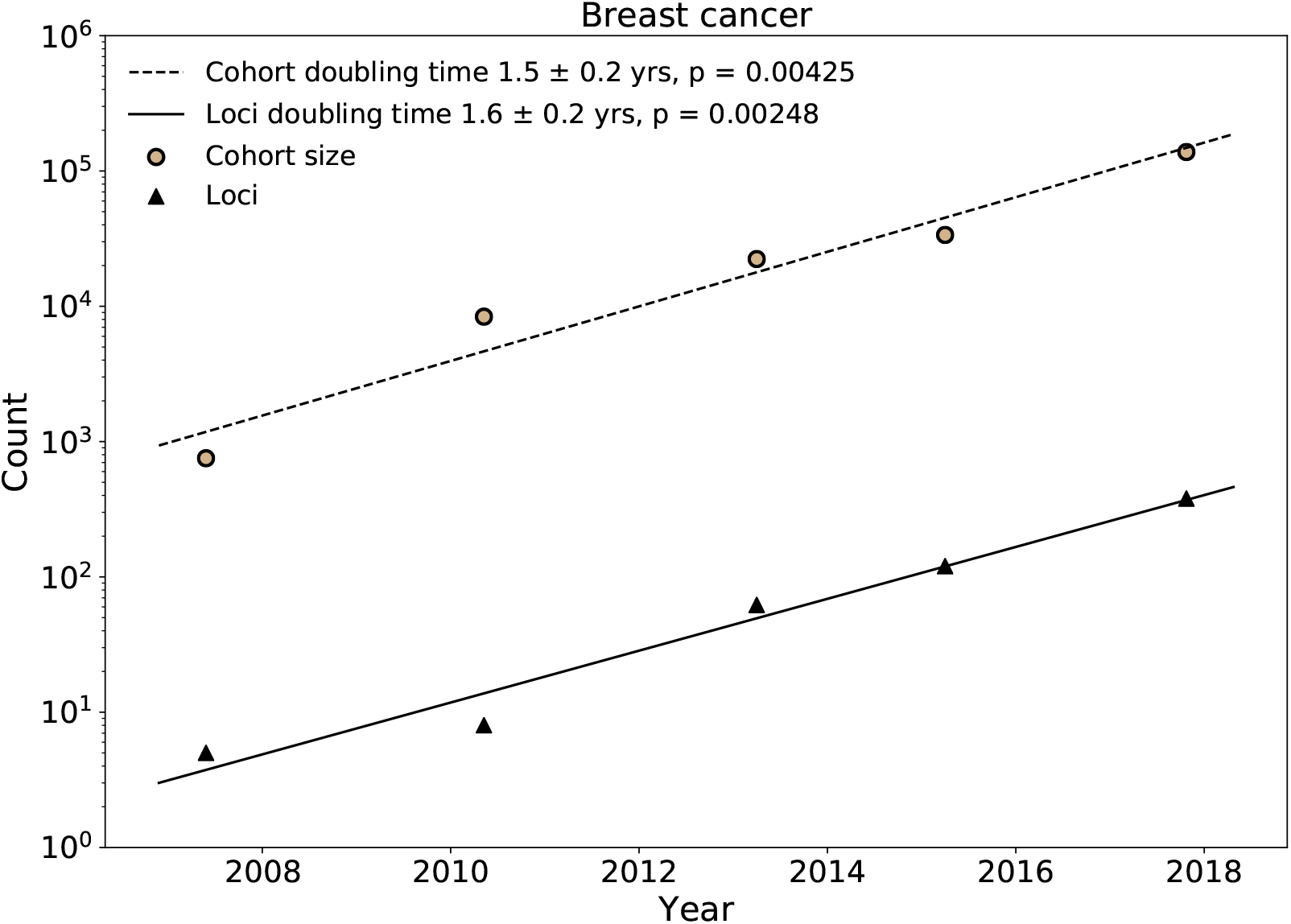
Growth rate of sample size and number of loci for GWAS investigating breast cancer. The scatter plot shows that both the sample size and number of loci increase through time for GWAS investigating breast cancer. The cohort size and numbers of loci increase linearly in log scales during last decade. X-axis indicates time and y-axis indicates the counts. Every points is a single study recorded in GWAS.

Analogous results for a well-studied quantitative trait, blood triglyceride levels, show similar progress from 2010 to 2020 (Fig. 3). As of 2010, the largest study had a cohort size of 96,598, and 65 loci had been identified. As of 2020, the largest cohort was 283,251, and 452 loci had been identified. The doubling time for cohorts was estimated at *τ_N_* = 1.8 ± 0.5 years, and the doubling for loci was estimated at *τ_L_* = 1.8 ± 0.2 years (Fig. 4).

**Fig 3.**
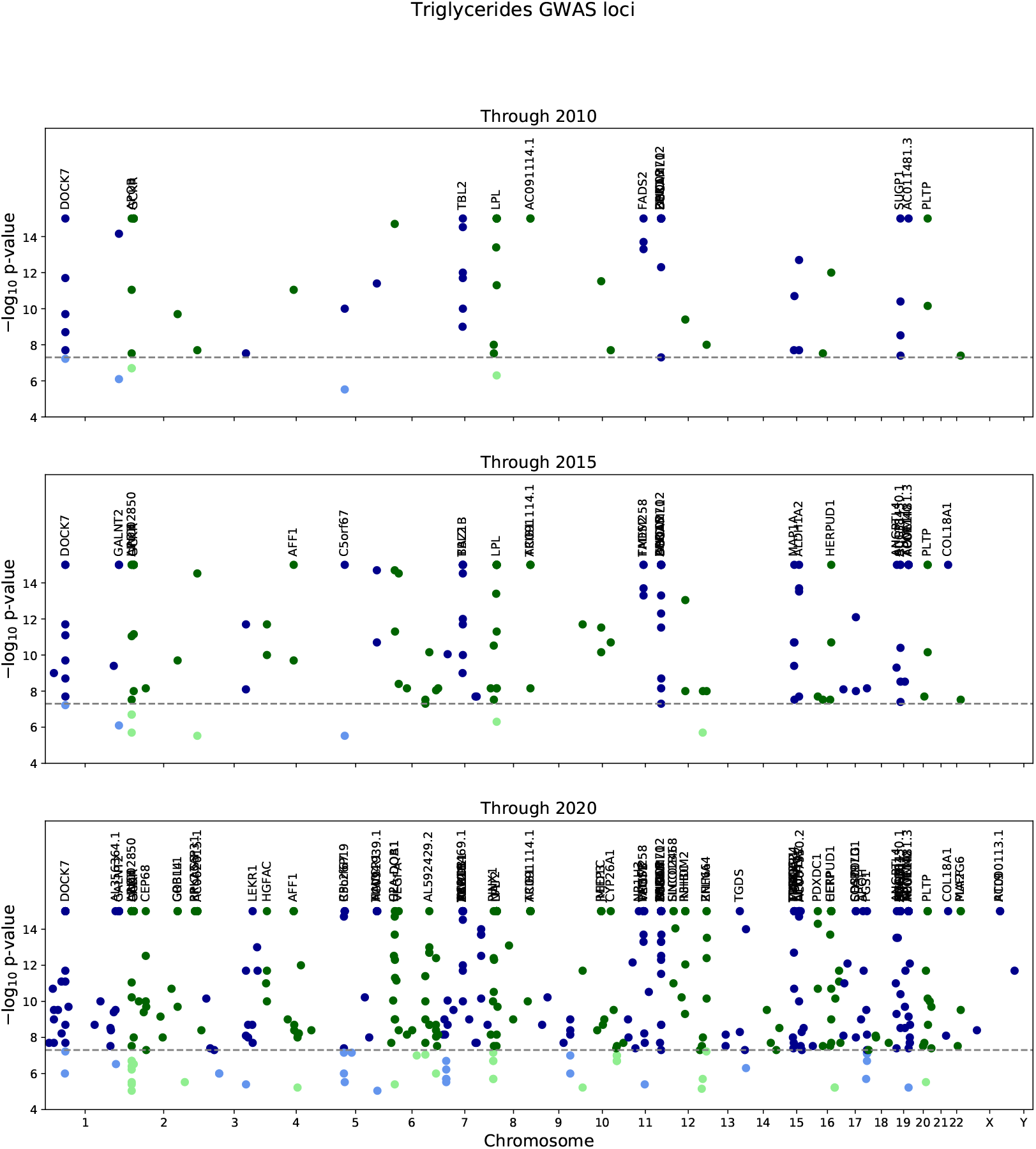
GWAS progress for triglyceride cohorts and loci. Manhattan plots depict GWAS findings for blood triglyceride levels as of 2010 (top panel), 2015 (middle panel), and 2020 (bottom panel). In each panel, the x-axis represents genomic coordinates to scale, and the y-axis is the −log_10_ p-value for a GWAS association with a SNP; a dashed line indicates the genome-wide significance threshold, *p* = 5 × 10^−8^. The SNP color alternates blue/green by chromosome, with lighter colors for findings below threshold and saturated colors above threshold.

**Fig 4.**
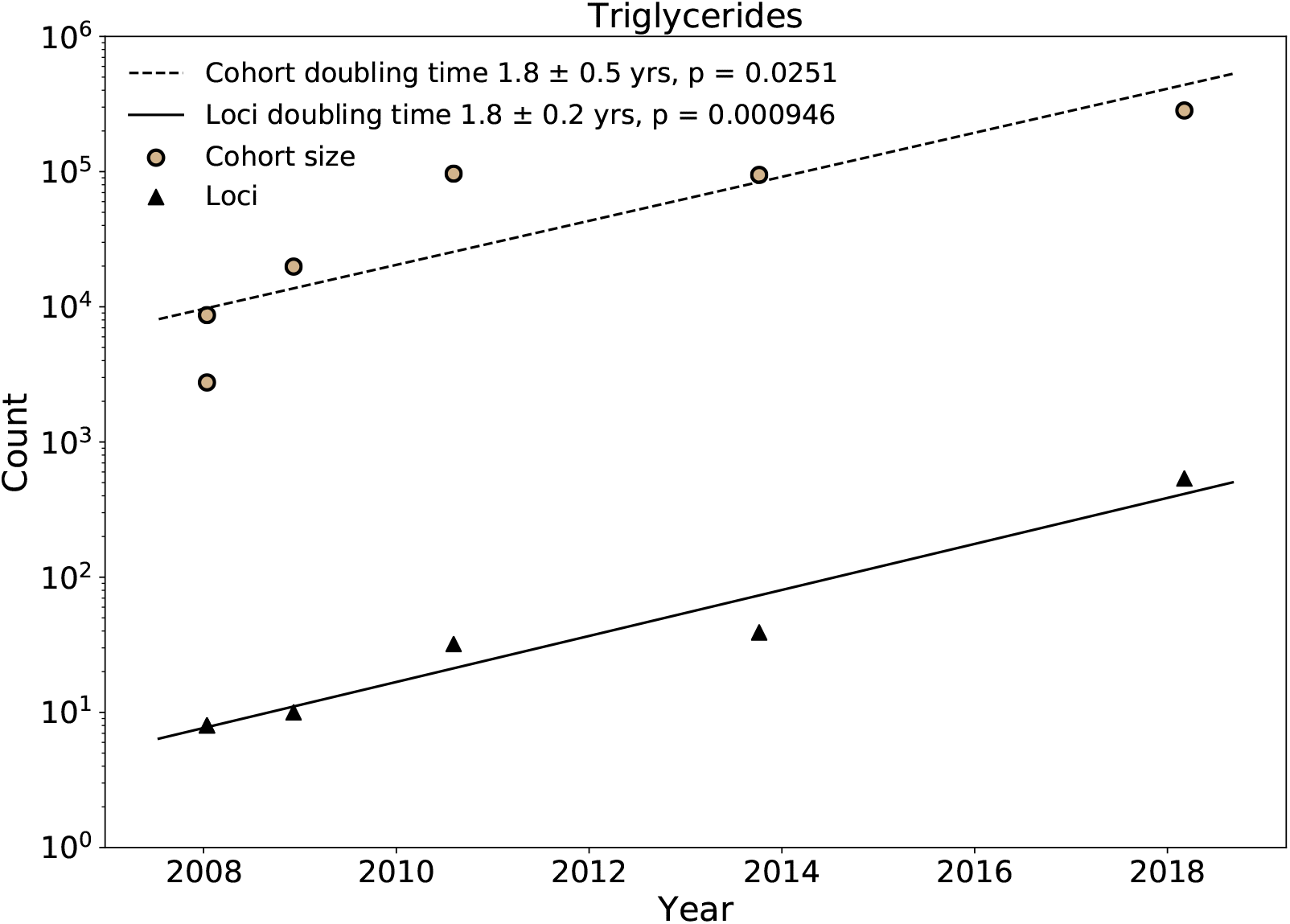
Growth rate of sample size and number of loci for GWAS investigating triglycerides in blood. The scatter plot shows that both the sample size and number of loci increase through time for GWAS investigating triglycerides in blood. The cohort size and numbers of loci increase linearly in log scales during last decade. X-axis indicates time and y-axis indicates the counts. Every points is a single study recorded in GWAS.

### Implications for effect size distributions

Results for disease traits where sufficient studies have been reported to estimate doubling times show a general agreement between the doubling time for cohort size and the doubling time for the number of significant loci (Fig. 5). For 23 of the 33 traits analyzed, the null hypothesis of equal doubling times for cohort size and loci cannot be rejected at p = 0.05.

**Fig 5.**
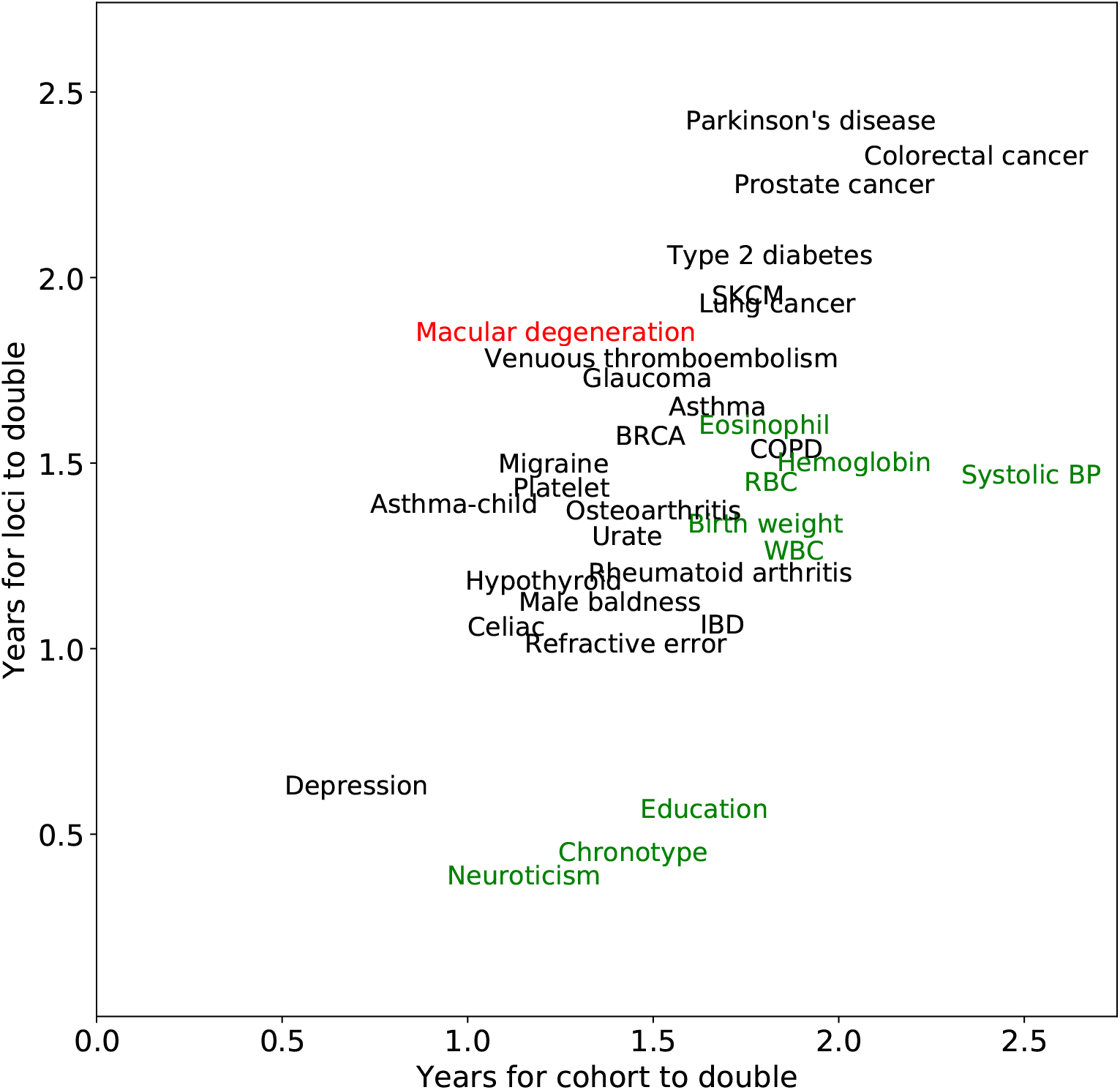
Comparison between years of doubling for loci and cohort in GWAS studies. Each term denotes a trait recorded in GWAS catalog. The corresponding x-axis indicates the cohort doubling time for studies investigating that trait, and y-axis indicates the captured loci doubling time. Terms in black shows that the doubling time for study cohort and loci is approximately the same. Traits that have a significant larger doubling time for cohort than loci are marked in green. Traits with significant larger doubling time for loci are marked in red.

The only trait for which the number of loci is doubling more slowly than the cohort size is age-related macular degeneration, with a cohort doubling time of 1.2 years and a loci doubling time of 1.8 years. Early studies of this phenotype with small cohorts nevertheless discovered significant loci with large effects. This may explain the subsequent slower rate of discovery.

Traits for which loci are doubling faster than cohort size include neuroticism, chronotype, and education, with cohorts doubling every 1.2 to 1.6 years and loci doubling every 0.4 to 0.6 years. An important aspect noted in a recent educational attainment study is the difficulty in controlling for environmental effects that are correlated with causal variants and overstate the causal effect [15].

Well-defined doubling times imply a power-law for the cumulative distribution *L*(*R*^2^) of the number of loci with effect size *R*^2^ or greater,

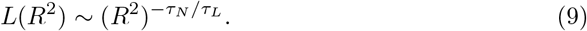

Equivalence in doubling times leads implies the specific exponent of −1,

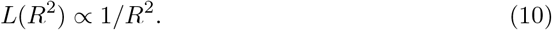

Algebraic or power-law decay is synonymous with a scale-free distribution and is also known as Zipf’s law. This conjecture is generally consistent with the omnigenic view of complex traits.

### Oracular hard priors

The previous results describe why increasing cohort sizes have increasing power to detect additional GWAS loci: the cumulative effect size distribution for many traits has a power-law functional form approximately proportional to 1/*R*^2^, and a proportional increase in the cohort size *N* can reveal the same proportional increase in genome-wide significant loci.

In this section, we hold the population size fixed and instead present results for oracular hard priors. The term “hard” indicates a frequentist framework in which only hypotheses within a pre-specified subset are tested, and a multiple testing correction is applied according to the number of hypotheses within this subset. The alternative “soft” prior in a Bayesian framework would be a prior probability assigned to each hypothesis, for example arising from predictions of functional impact. The hard prior is valuable in permitting an analytical treatment for general insight into the value of priors.

The “oracular” property indicates that all the hypotheses that in reality fall under the alternative hypothesis are guaranteed to be within the subset selected for testing. This again is a simplification that overestimates any real-world prior, which invariably will create false negatives by not testing some of the hypotheses that really fall under the alternative.

Fewer hypotheses tested corresponds to a reduced multiple-testing burden. This in turn implies a less stringent significance threshold and a greater power to detect positives within the prior region.

Studies of cancer somatic mutations to identify cancer drivers (rather than cancer susceptibility loci) regularly apply hard priors to reduce the testing burden from passenger mutations (Fig. 6). The universe of all possible hypothesis tests includes every somatic mutation identified in a tumor cell versus the germ line, regardless the fraction of tumor cells that carry the mutation or the predicted functional impact of the mutation. Loss-of-function mutations, including deletions, frameshifts, and nonsense mutations, are generally prioritized for testing. Tumor suppressor genes often have these types of mutations. Missense mutations that arise recurrently in independent individuals are also prioritized for testing, with a gain-of-function hypothesis. Oncogenes often have recurrent non-synonymous mutations, or recurrent loss of specific regulatory domains leading to constitutive activity. Mutations that are less likely to have functional consequences, including non-recurrent non-synonymous mutations and mutations in non-protein-coding regions, are often not tested at all. The number of neutral mutations can be orders of magnitude larger than the number of driver mutations [16, 17] Consequently only a small fraction of the observed somatic mutations are tested, considerably reducing the multiple testing burden.

**Fig 6.**
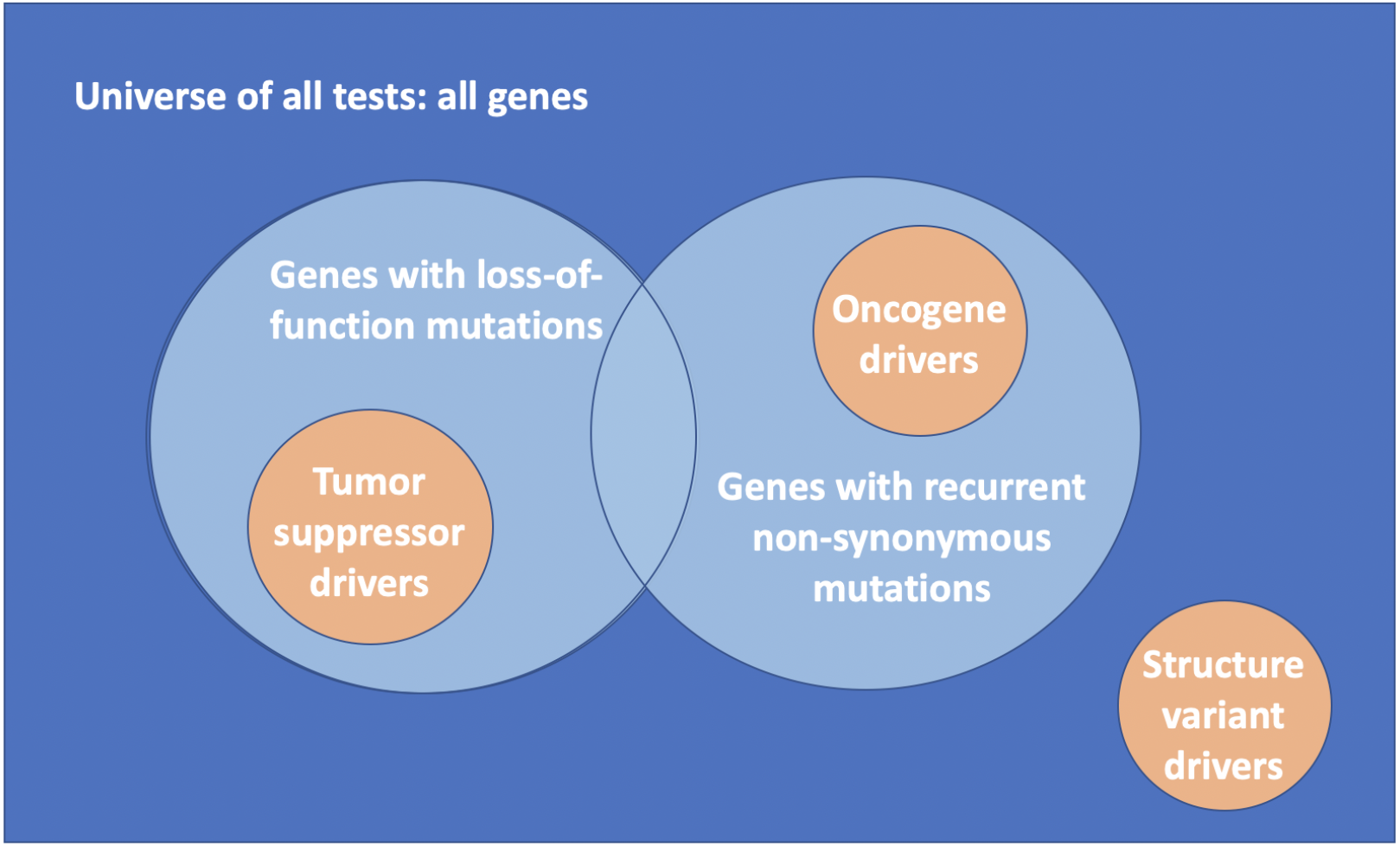
Hard priors in the context of cancer somatic mutations. The universe of all tests (dark blue) includes all identified somatic mutations, regardless of frequency within cancer cells or recurrence across individuals. Hard priors often restrict tests to loss-of-function mutations (light blue) characteristic of tumor suppressors (orange) and to recurrent non-synonymous mutations (light blue) characteristic of oncogenes (orange). These priors may exclude unanticipated classes of driver mutations, for example structural variants (orange) in which copy-number amplifications lead to changes in gene activities that drive cancer. While oracular hard priors have the property that all true findings are within the subset selected for testing, in real-world applications true findings will fall outside the prior and can increase the false-negative rate.

Hard priors for cancer studies are easier to formulate, given the strong functional effects assumed for cancer driver mutations. Similarly, positional cloning studies to identify Mendelian disease loci successfully rely on sequencing to identify germ line mutations with strong functional effects and high penetrance. For GWAS of complex disorders, genetic variants predisposing to disease are less readily identified by strong effects on protein structure or function. Nevertheless, priors that focus attention on genes expressed in a tissue of interest at an appropriate developmental stage, or priors that focus attention on SNPs that are within gene-based boundaries or coincide with regulatory regions, have been discussed and applied.

For a general mathematical framework, we define the strength *S* of a hard prior as the fold-reduction in the number of hypotheses tested: rather than testing all *A* hypotheses, only a subset of size *A/S* is tested. The stronger the prior, the larger the value of *S*, and the smaller the multiple testing burden. For a GWAS, a p-value cutoff of 5 × 10^−8^ is usually used for genome-wide tests. A prior with strength *S* = 100, roughly equivalent to testing SNPs in 250 rather 25,000 genes, would change the p-value cutoff to 5 × 10^−6^.

This type of approach can identify some variants that are not identified using a conventional threshold. For example, a breast cancer susceptibility GWAS investigating breast cancer risk identified the SNP rs11571833 in BRCA2 with p-value of 2 × 10^−6^. The BRCA2 gene is a known tumor suppressor contributing to DNA repair. Based on this prior knowledge, the SNP is likely to be a true finding. A strong prior, for example a prior limiting tests to SNPs close to known breast cancer risk factors, would yield a significant finding for rs11571833, which could not be captured as a significant hit with the normal p-value cutoff for GWAS studies. This type of prior could suffer from false negatives, however, in excluding most of the genome from testing.

Instead of devising and applying priors, an alternative strategy would be to accumulate larger cohorts for greater power to detect real effects. Larger cohorts and improved priors could of course be pursued simultaneously. Nevertheless, given the 18–24 month doubling time for cohort size and the similar amount of time required to develop and benchmark a new computational method, it is worthwhile to consider the prior strength *S* required to give the same boost in power as simply waiting one to two years for a larger study.

### Power as a function of prior strength and cohort size

Consider two studies with equal power, represented by the normal quantile *z_II_* for the false-negative rate for a particular effect size. Typically a power of 80% is desired for a small effect, with *z_II_* = −0.84. One study has a larger population of size *N*_1_ and applies no prior, *S*_1_ = 1. This study has a full multiple testing burden, represented by the normal quantile *ζ*. For GWAS, *ζ* corresponds to the quantile for a two-sided p-value of 5 × 10^−8^, or *ζ* = 5.45. As shown in the methods using a steepest descents approximation for the normal distribution survival function (or the complementary error function),

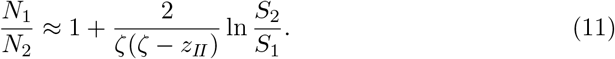

Using the values described above for GWAS, *ζ*(*ζ* – *z_!I_*)/2 = 17.2. The logarithmic term divided by this value is small, permitting the approximation 1 + *ϵ* ≈ *e^ϵ^* with error *ϵ*^2^.

Using this approximation,

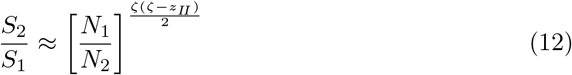

The population size is exponentially more important than the prior in determining the critical effect size for a study, defined as the effect size *R*^2^ that can be detected with 80% power at significance threshold *αS*, where *α* is the threshold without a prior and *S* is the prior strength (Fig. 7). The contour lines of equal power as a function of prior strength and population size have the steep slope of about 17 on this numerically exact log-log plot, exactly as predicted by the analytical approximation.

**Fig 7.**
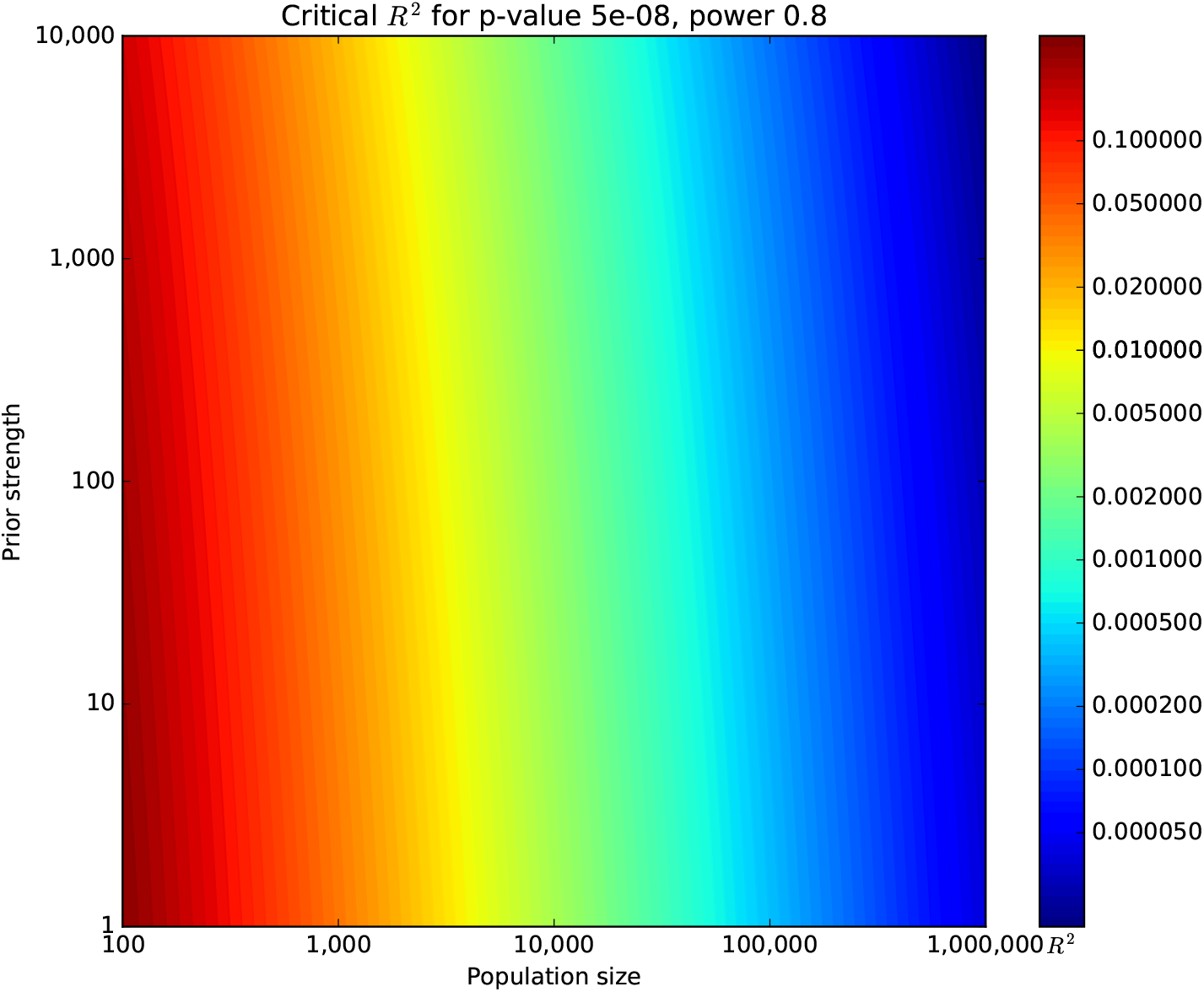
Critical *R*^2^ for *p* = 5 × 10^−8^ at power = 0.8 as a function of prior strength and population size. Colors indicate contour lines of equal power to detect an effect in the context of GWAS. Power changes rapidly left-to-right, reflecting the strong dependence of power on population size. Power changes very slowly bottom-to-top, reflecting the weak ability of priors to boost power.

The full contour plot of critical *R*^2^ is summarized by a numerically exact calibration curve depicting the prior strength required to match the power increase from a larger cohort (Fig. 8). A 1.2-fold larger population could be matched by a prior of strength 16.8, roughly equivalent to restricting tests to genes specific to a tissue of interest. A 1.4-fold larger population could be matched by a prior strength 124.2, roughly equivalent to restricting tests to a pathway of about 100-200 genes. Doubling the cohort size would require a prior strength 4321.5. This would almost certainly violate the oracular property by restricting tests to about 1–10 genes, or to about 500 of the effectively 2 million independent SNPs usually assumed for the GWAS multiple testing burden. This type of prior can be useful for validation studies but would have an unacceptably larger false-negative rate for a discovery study.

**Fig 8.**
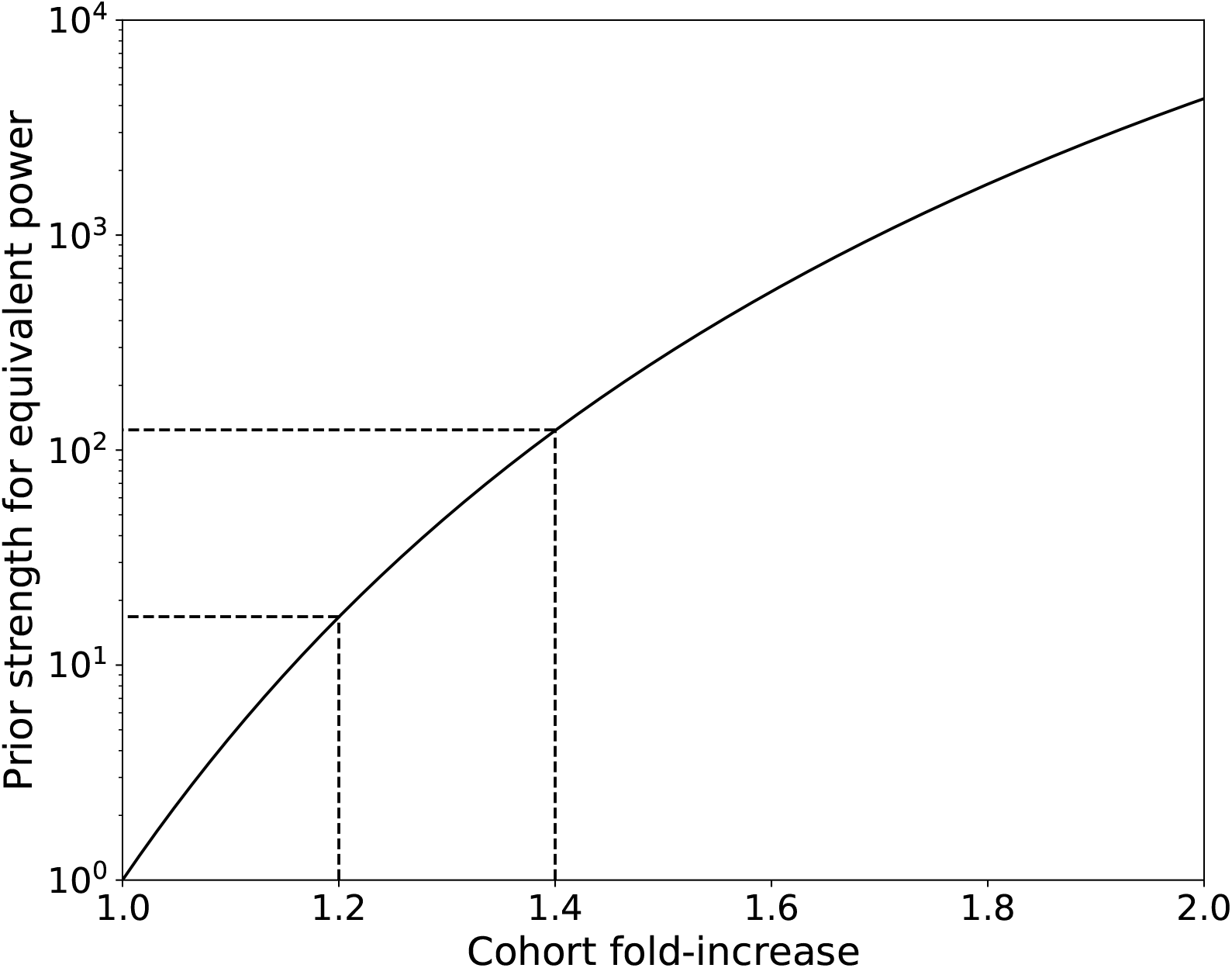
Calibration curve for power and population size. Numerically exact results are shown for prior strength *S* required to achieve the same power as a simple fold-increase in population size with no prior.

## Discussion and Conclusion

Despite long-standing efforts to exploit prior knowledge to boost the power of GWAS, RNA-Seq, and other genome-wide tests, including our own efforts to develop and apply gene-based priors for GWAS [8, 9], traditional unbiased univariate tests with a Bonferroni multiple-testing correction are essentially the only method in use. One hypothesis is that genetics researchers performing GWAS do not appreciate the power gains that could be had by using the most recent methods. An alternative hypothesis supported by our findings is that effort is better spent collecting larger cohorts, performing meta-analyses, and avoiding the risk that priors that attempt to increase power will actually degrade performance by increasing the false-negative rate. Supporting the alternative hypothesis is are no-free-lunch theorems that prove that priors that improve performance for one class of problems must degrade performance for other classes [18].

What, then, is the value of priors in GWAS? We are ourselves convinced that priors that attempt to boost statistical significance are not a productive area for future research. We do think, however, that they have an important role in bridging between statistical significance—which SNP-based associations will replicate in a validation cohort—and biological mechanism. While larger cohorts will provide stronger statistical associations, they will not necessarily identify which SNP or SNPs in a linkage disequilibrium block is most likely to be responsible for the observed effect, or even how many independent effects are contributed by a linkage region. Similarly, purely statistical approaches based on GWAS data do not necessarily link the causal SNPs to the gene or genes whose function they affect. These types of problems are an excellent opportunity for the use of priors, whether as part of a hypothesis-testing framework or using Bayesian methods for inference over multiple types of data and existing biomedical knowledge.

## Acknowledgements

JSB and DEA acknowledge funding from US NIH R01HL141989. JSB acknowledges funding from US NIH U01CA217846.

## Author Contributions

All authors designed the study, analyzed the results, and drafted the manuscript. JC and JSB implemented the methods and performed the mathematical and computational analyses.

